# The Nuclear RNAi Pathway Regulates DAF-16/FOXO to Control *C. elegans* Longevity and *Dauer* Entry

**DOI:** 10.1101/2022.10.21.513151

**Authors:** Sweta Sarmah, Evandro A. De-Souza, Silas Pinto, Adam Antebi, Marcelo A. Mori

## Abstract

*Caenorhabditis elegans* with impaired insulin/IGF-1 receptor signalling (IIS) or with germline ablation live longer and this phenomenon is entirely dependent on the transcription factor DAF-16 - the *C. elegans* homolog of the class O of the forkhead box transcription factors (FoxO). In a candidate RNAi screen designed to search for new modifiers of DAF-16 function among genes involved in histone modification and/or small RNA-mediated silencing pathways, we found *nrde-1, wago-1*, and *adr-1* as positive regulators of DAF-16. We confirmed by several methods and in different models that DAF-16 translocation to the nucleus and, subsequently, its function is tightly controlled by these genes and narrowed down to components of the NRDE complex and the nuclear RNAi pathway as key DAF-16 modulators. Importantly, we found that the NRDE pathway controls DAF-16-mediated longevity and *dauer* entry. Our epistasis data indicate that *nrde-1* interacts with *akt-1* to control lifespan. We also demonstrated that NRDE-1 acts downstream of AGE-1/PI3K and partially requires mTORC2 and AKT-1 to control DAF-16 translocation. These results unveil a mechanism of regulation of *dauer* formation and longevity in *C. elegans* via nuclear RNAi-mediated modulation of DAF-16 function in a manner that involves the mTORC2-AKT axis.

## INTRODUCTION

Aging comes with several changes such as physiological, psychological, social, and many others, leading to increased vulnerability to different diseases such as type 2 diabetes, cancer, hypertension, cardiovascular diseases, neurodegenerative disorders, (Partridge et al. 2018) and ultimately death. The first demonstration that aging can be modified genetically was conducted using the free-living nematode *C. elegans*, in which loss-of-function mutations in the insulin/IGF-1 signalling (IIS) pathway resulted in the extension of the worm lifespan (Kenyon et al. 1993; Friedman & Johnson 1988; Kimura et al. 1997a). These seminal discoveries in worms paved the way for subsequent studies that dissected the involvement of the IIS and other pathways in aging regulation across the evolutionary spectrum (Kenyon 2010). Numerous interventions that extend the lifespan in this model also impact animals’ stress resilience, metabolism, and fertility (Hansen et al. 2013). Accordingly, it was observed that the elimination of germ cells extends lifespan in invertebrate species, such as flies and worms (Flatt et al. 2008; Hsin & Kenyon 1999), by the activation of a variety of stress response pathways in somatic tissues (Steinbaugh et al. 2015a; Hsin & Kenyon 1999; Antebi 2013).

The class O of the forkhead transcription factors (FoxO) is necessary for both IIS deficiency- and germline ablation-induced longevity (Kenyon et al. 1993; Hsin & Kenyon 1999). DAF-16 is the sole FoxO homolog in *C. elegans*, has multiple isoforms (a, b, and d/f) and a high degree of homology to human FOXO3A (Ogg et al. 1997; Lin et al. 1997). DAF-16 was first identified in *C. elegans* as a gene required for *dauer* - a larval stage of diapause induced by unfavourable conditions (*e.g.*, high temperature, low food availability, high population density, and/or pheromones) which confers worms resistance to stress and dramatically extends worm lifespan (Golden & Riddle 1984; Golden & Riddle 1982). Like DAF-16, other components of the IIS pathway play a major role in controlling the *dauer* formation (Vowels & Thomas 1992; Kimura et al. 1997b). Loss-of-function mutations in genes encoding the IIS components DAF-2 (insulin/IGF-1 receptor ortholog), AGE-1 [part of the phosphoinositide-3 kinase (PI3K) complex], PDK-1 (3-phosphoinositide dependent protein kinase 1 ortholog) or AKT-1 and −2 (AKT serine/threonine kinases 1 and 2 orthologs) can result in constitutive *dauer* phenotype in a way that interacts with DAF-16 (Ogg & Ruvkun 1998; Paradis & Ruvkun 1998a; Paradis et al. 1999a). This is because a large extent to which IIS controls transcription of genes involved in growth, metabolism, stress resistance, apoptosis, and cell cycle is through suppression of DAF-16 function (Murphy et al. 2003a).

When DAF-2 signalling is triggered, it leads to the activation of a downstream kinase cascade starting with AGE-1 (Dorman et al. 1995). AGE-1 activation increases PI(3, 4, 5)P3/PI(4, 5)P2 ratio, thus inducing PDK-1 (Paradis et al. 1999b). In humans, PDK1 phosphorylates AKT1 at Thr-308 and, in a parallel signalling cascade, activation of mTOR Complex 2 (mTORC2) phosphorylates AKT1 at Ser-473 (Alessi et al. 1996; Sarbassov et al. 2005). These residues are present in *C. elegans* AKT-1 too, and a similar level of regulation of the IIS pathway by PDK-1 is observed in worms (Hertweck et al. 2004; Paradis et al. 1999b). Upon phosphorylation, AKT is activated and phosphorylates DAF-16, blocking its nuclear entry and resulting in sequestration in the cytosol and inhibition of its transcriptional activity (Paradis & Ruvkun 1998b; Paradis et al. 1999b). Therefore, defects in upstream components of the IIS reduce DAF-16 phosphorylation, resulting in its nuclear translocation and subsequent transcriptional activation of DAF-16 target genes, which in turn allow worms to be more stress-resistant and have a longer lifespan (Murphy et al. 2003b).

As previously mentioned, worms that have their germline cells genetically or physically (with lasers) ablated, are long-lived, and the longevity phenotype is manifested in a DAF-16-dependent manner (Hsin & Kenyon 1999). Strong nuclear DAF-16 localization is also observed in animals lacking the germline, and like in the case of *daf-2* mutants, longevity and enhanced DAF-16 nuclear translocation in germline-less worms are suppressed by removing DAF-18/PTEN – a phosphatase that antagonizes AGE-1/PI3K (Berman & Kenyon 2006). Although both the inhibition of the IIS and the ablation of germ cells extend lifespan through DAF-16, *daf-2* mutants display an even longer lifespan when it has their gonads ablated by laser (Arantes-Oliveira et al. 2003), suggesting that non-overlapping mechanisms act to control longevity by the IIS and germline pathways. Accordingly, it was observed that distinct mechanisms are involved in the regulation of DAF-16 by these two pathways. For instance, *daf-9* and *daf-12* were observed to be necessary for DAF-16 nuclear localization and longevity in germline-less animals, but are not necessary for *daf-2* mutants (Berman & Kenyon 2006). DAF-9 is involved in the synthesis of the signalling hormone molecule (*i.e.* dafachronic acid), which activates the nuclear hormone receptor DAF-12 (Motola et al. 2006). A possibility previously raised is that DAF-12 is directly regulating DAF-16 nuclear translocation (Lapierre & Hansen 2012; Dowell et al. 2003). TCER-1, a conserved transcription elongation factor, was also shown to exclusively regulate DAF-16 nuclear localization in germline-ablated animals (Ghazi et al. 2009). Thus, IIS and germline signalling pathways have both overlapping and exclusive components that regulate DAF-16 function and longevity.

In addition to the regulation by transcriptional factors such as DAF-16, other players, such as chromatin-regulating proteins are essential for enabling cells to coordinate gene expression during stress and aging. Accordingly, it was observed that epigenetic factors play an essential role in lifespan extension in the majority of anti-aging interventions tested in *C. elegans*, including in IIS and germline signalling pathways (Denzel et al. 2019). For example, it was observed that microRNAs [small RNAs involved in gene expression regulation, whose activity and expression decrease with aging (Mori et al. 2012; Kato et al. 2011)] play a role in regulating DAF-16 activity in germline-ablated animals (Boulias & Horvitz 2012). SIR-2.1 – a histone deacetylase – physically interacts with DAF-16 during heat stress, and is partially required for the lifespan increase observed in *daf-2* mutants (Berdichevsky et al. 2006). Finally, other epigenetic regulators, such as the histone demethylase JMJD-3.1 and the chromatin remodeller LET-418 have been shown to participate in the observed increase in longevity of germline-ablated animals (Labbadia & Morimoto 2015; De Vaux et al. 2013), but it is still elusive if they do that by regulating DAF-16 nuclear localization.

To try to better understand how epigenetic mechanisms influence DAF-16 function in transcription and longevity, we undertook an RNAi mini-screen of genes involved in chromatin regulation and small RNA pathways, monitoring DAF-16 nuclear localization as a readout of its activity. We were able to identify that *nrde-1*, *wago-1*, and *adr-1* regulate DAF-16 nuclear localization and function in different contexts, including in animals lacking the germline or after heat shock. We also found that *nrde-1*, *wago-1*, and *adr-1* play a role in *dauer* entry and lifespan extension in worms deficient for the IIS pathway, demonstrating that the control of DAF-16 by these genes impacts worm physiology. Finally, we observed that all members of the NRDE family participate in DAF-16 regulation to some extent and the mechanism through which the nuclear RNAi pathway controls DAF-16 nuclear translocation partially involves mTORC2 and AKT-1.

## RESULTS

### Downregulation of *nrde-1, wago-1*, or *adr-1* inhibits DAF-16 translocation

To search for new regulators of DAF-16 function among genes involved in histone modification and/or small RNA-mediated silencing pathways, we performed a candidate RNAi screen using the clones listed in Table S1 and assessed GFP::DAF-16 nuclear localization (*muIs109*) in a long-lived strain that lacks the germline, namely *glp-1(e2141)* mutant. Briefly, L1 worms were exposed to RNAi and nuclear vs. cytoplasmic GFP localization was scored on day 1 of adulthood. Control worms, *i.e., glp-1(e2141)* mutant worms treated with RNAi against luciferase (De-Souza et al. 2019), were used to set the reference for the condition with the maximal number of intestinal cells displaying nuclear DAF-16 localization. This screening strategy limited us to the identification of positive regulators of DAF-16 translocation in germline-less worms. Among 36 tested clones, we found 4 that resulted in inhibition of DAF-16 nuclear localization in 3 independent experiments (Table S1). These clones targeted genes encoding NRDE-1 [a protein involved in nuclear RNAi – (Burkhart et al. 2011)], WAGO-1 [an argonaute involved in the endogenous siRNA pathway - (Gu et al. 2009a)], ADR-1 [a protein involved in A-to-I RNA editing - (Knight & Bass 2002)], and MES-4 [a histone methyltransferase - (Bender et al. 2006)]. Other clones inhibited DAF-16 nuclear localization in 2 out of 3 replicates (*i.e.*, *pgl-1* and *set-25*) or in 1 out of 3 replicates (*i.e.*, *hda-1* and *ire-1*).

To validate whether these genes might participate in the regulation of DAF-16 function, we tested whether RNAi against some of these genes could also downregulate a DAF-16 target gene reporter (*i.e.*, *sod-3p::GFP/muIs84*). *nrde-1, wago-1*, *adr-1, hda-1*, and *set-25* RNAi reduced the expression of the *sod-3* reporter construct by at least some level (Figure S1A and Table S2). Furthermore, we tested whether the same genes affected SKN-1 nuclear localization. SKN-1 is yet another stress response transcription factor that is required for the longevity phenotype of germline-less worms (Wei & Kenyon 2016; Steinbaugh et al. 2015b). Interestingly, *adr-1, hda-1*, and *wago-1* RNAi also reduced SKN-1 nuclear translocation, while *nrde-1* and *set-25* RNAi did not (Table S3). Considering that *nrde-1, wago-1*, and *adr-1* RNAi were the only clones that consistently inhibited DAF-16 translocation as well as DAF-16 target gene reporter expression, we decided to focus on these genes and explore their mechanisms of regulation of DAF-16.

To test whether *nrde-1, wago-1*, and *adr-1* could also control DAF-16 nuclear localization induced by other types of stimuli, we silenced these genes and measured the translocation of diverse DAF-16::GFP reporter transgenes in heat-shocked worms (Figure 1A-C) or *daf-2(e1370)* mutant worms (Figure 1D). We found inhibition of DAF-16 nuclear localization across all the models when we silenced *nrde-1, wago-1*, or *adr-1*, being *nrde-1* RNAi the one that elicited the strongest inhibition. This was observed regardless of the stimulus for DAF-16 translocation (*i.e.*, lack of germline, *daf-2* loss of function, or heat stress) and happened even when DAF-16::GFP was expressed using an independent promoter (i.e., *ges-1p*) or in the absence of its *daf-16b* isoform. Together, these results point to the involvement of *nrde-1, wago-1*, and *adr-1* in post-transcriptional regulation of DAF-16 nuclear localization by several stimuli.

**Figure 1.**
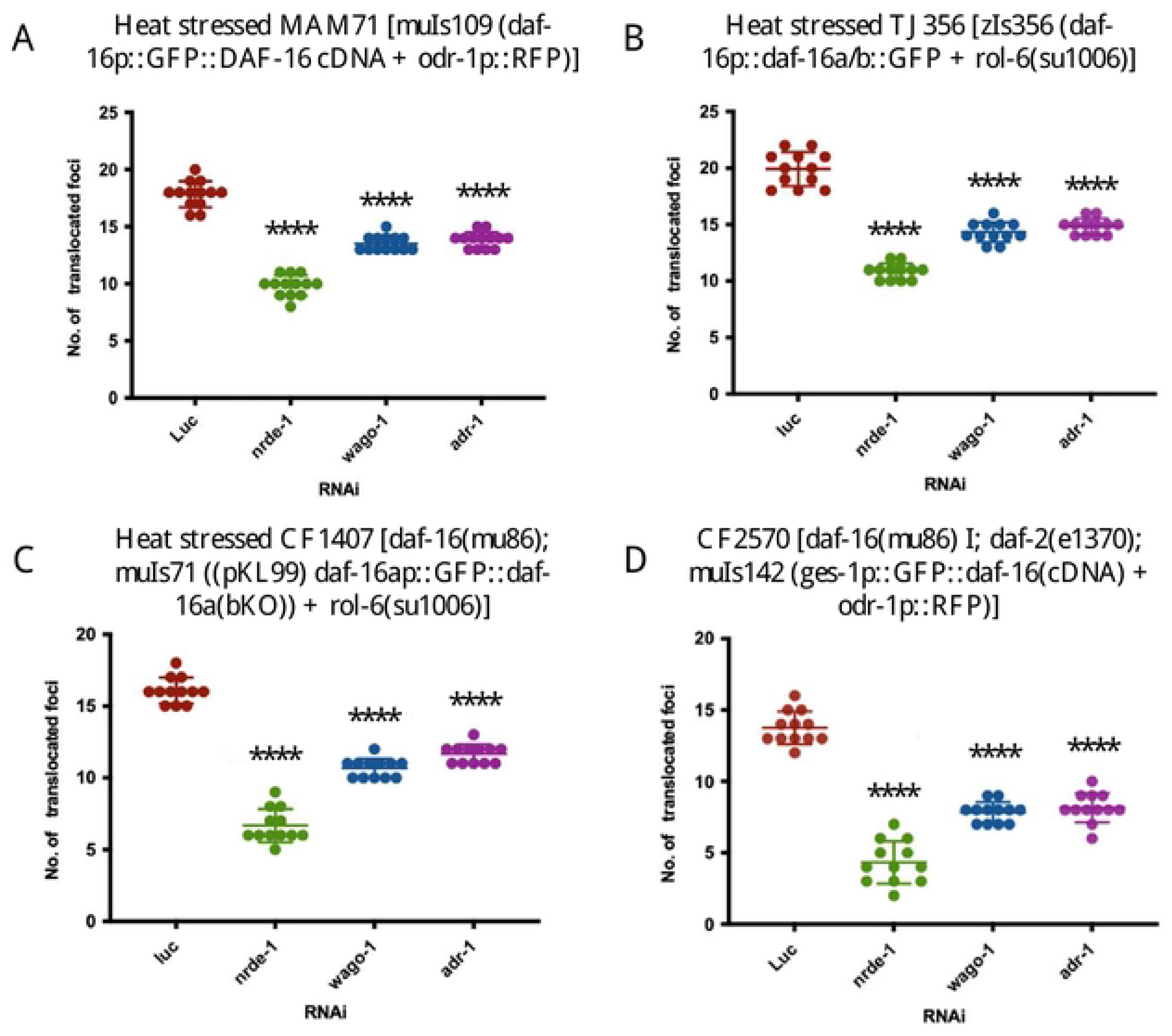
DAF-16 translocation is inhibited by *nrde-1*, *wago-1* and *adr-1* RNAi. *nrde-1, wago-1*, and *adr-1* were silenced by RNAi and RNAi targeting luciferase (luc) was used as a control. The number of DAF-16::GFP positive nucleus (the translocation of DAF-16 from the cytoplasm to the nucleus) was scored in microscopic images obtained using Cytation 5. A, MAM71 (n = 13 per group). B, TJ356 (n = 12 per group). C, CF1407 (n = 12 per group). D, CF2570 (n = 12 per group). These experiments were repeated 3 times with similar results. In A-C, worms were maintained at 20 °C and heat-shocked at 34 °C for 1.5 hours on the first day of adulthood. In D, worms were maintained at 15 °C till L2 and then transferred to 25 °C until adulthood. \*\*\*\**P* < 0.0001 vs. luc. Two-tailed student *t*-test was used for individual comparisons against the control group (luc).

### Downregulation of *nrde-1*, but not *wago-1* or *adr-1* inhibits DAF-16 expression

We then sought to quantify the intensity of GFP as a measurement of the overall level of expression of DAF-16. A decrease in DAF-16 levels (*i.e.*, GFP) was observed in the transgenic models upon silencing with *nrde-1* RNAi, but not with the other RNAi clones (Figure S1B-E). However, this decrease was not as pronounced as the decrease in DAF-16 nuclear localization, suggesting that NRDE-1 controls DAF-16 both at the expression as well as the translocation level, while WAGO-1 and ADR-1 are primary regulators of DAF-16 translocation. Interestingly, DAF-16 levels also decreased upon *nrde-1* RNAi in the CF2470 strain which expresses *GFP::daf-16(cDNA)* under the control of the *ges-1* promoter, indicating a possible post-transcriptional regulation.

### Downregulation of *nrde-1, wago-1*, or *adr-1* reduces DAF-16 target gene expression

To assess the endogenous transcription activity of DAF-16 and confirm its interaction with *nrde-1, wago-1*, and *adr-1* in a transgene-free model, we used RT-qPCR to measure the DAF-16 target genes *sod-3, mdl-1*, and *scl-1* along with the isoforms of *daf-16a* and *daf-16b* in wild type worms (N2) subjected to either control RNAi or RNAi targeting *nrde-1, wago-1*, or *adr-1*. Worms were subjected to heat shock at 34°C for 1.5 hours on the first day of adulthood to induce DAF-16 function. Consistent with the hypothesis that endogenous DAF-16 is regulated by the candidate genes, we observed downregulation of all DAF-16 target genes and the *daf-16a/b* isoforms when *nrde-1* was silenced (Figure 2A-E). *daf-16a* and *daf-16b* were also reduced by *wago-1* and *adr-1* RNAi, but DAF-16 target genes only trended towards a statistically non-significant downregulation in these cases. These results agree with the observations using the transgenic worms, where the effects were much more pronounced when *nrde-1* was silenced.

**Figure 2.**
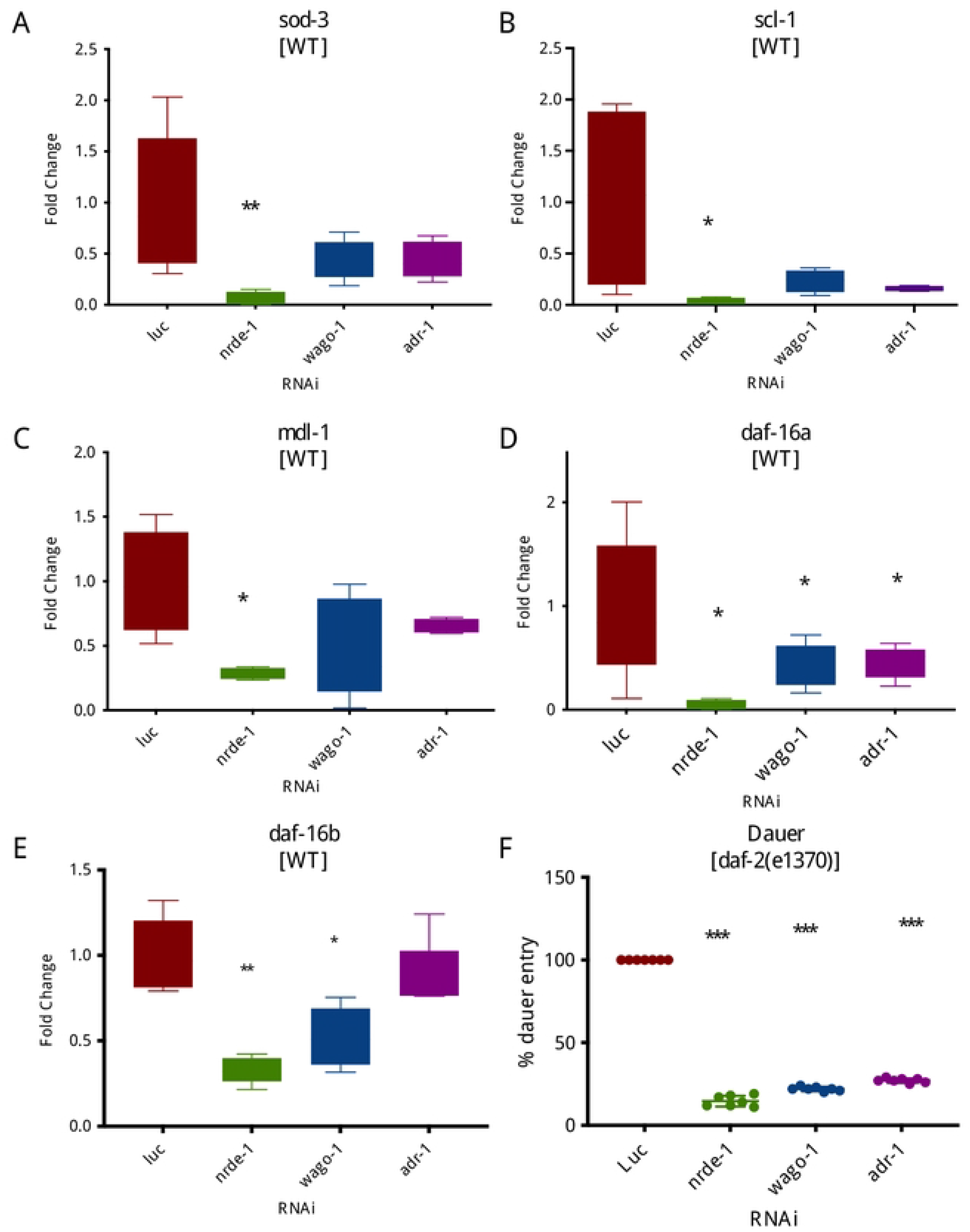
Endogenous DAF-16 function is inhibited by *nrde-1*, *wago-1* and *adr-1* RNAi. *nrde-1, wago-1*, and *adr-1* were silenced by RNAi and RNAi targeting luciferase (luc) was used as a control. A-E, Gene expression as measured by RT-qPCR in N2 (WT) worms (n = 5 pools of worms per group). (A) *sod-3*, (B) *scl-1*, (C) *mdl-1*, (D) *daf-16a*, and (E) *daf-16b.* These experiments were repeated 2 times with similar results. F, *Dauer* assay on *daf-2(e1370)* mutants (n = 100 worms per replicate in 7 replicates per group). These experiments were repeated 3 times with similar results. \**P* < 0.05; \*\**P* < 0.01; \*\*\**P* < 0.001. Two-tailed student *t*-test was used for comparison with the control.

### Downregulation of *nrde-1, wago-1*, or *adr-1* inhibits *dauer* formation

Another way to test DAF-16 endogenous function is by measuring *dauer* entry. *Dauer* is a resistant stage of nematode development and it can be promoted by mutations like in *daf-2.* To test the importance of *nrde-1, wago-1*, and *adr-1* on *dauer* formation, we silenced these genes in *daf-2(e1370)* mutants and quantified the number of progenies entering the *dauer* stage in the population. Silencing of the candidate genes resulted in a significant decrease in the *dauer* population (Figure 2F). Again, the effect was much stronger in the case of *nrde-1* silencing. Therefore, this result reveals that knocking down *nrde-1, wago-1*, or *adr-1* suppresses the constitutive *dauer* arrest phenotype of *daf-2(e1370)* mutant worms, which demonstrates that these genes are major regulators of endogenous DAF-16.

### Downregulation of *nrde-1, wago-1*, or *adr-1* reduces *C. elegans* lifespan

Once we found that *nrde-1, wago-1*, and *adr-1* affect endogenous DAF-16 function, we then decided to verify if they would be involved in lifespan regulation. To evaluate this, we measured the lifespan of wild-type (WT) and *daf-2(e1370)* mutant worms when *nrde-1, wago-1*, or *adr-1* was knocked down. In parallel, we also performed lifespan assays on DAF-16::GFP/*muIs109* transgenic worms and mutants for other members of the IIS pathway (Figure 3A), *i.e.*, *age-1(hx546)*, *pdk-1(mg142)*, and *akt-1(mg144)*. We observed a decline in the lifespan of WT and a strong suppression of longevity induced by *daf-2, age-1*, and *pdk-1* mutations when the candidate genes were silenced (Figure 3B-E). However, in *akt-1* mutants, *nrde-1* RNAi not only did not reduce lifespan but trended to slightly increase it (Figure 3F); demonstrating that *akt-1* is required for *nrde-1* RNAi to inhibit longevity.

**Figure 3.**
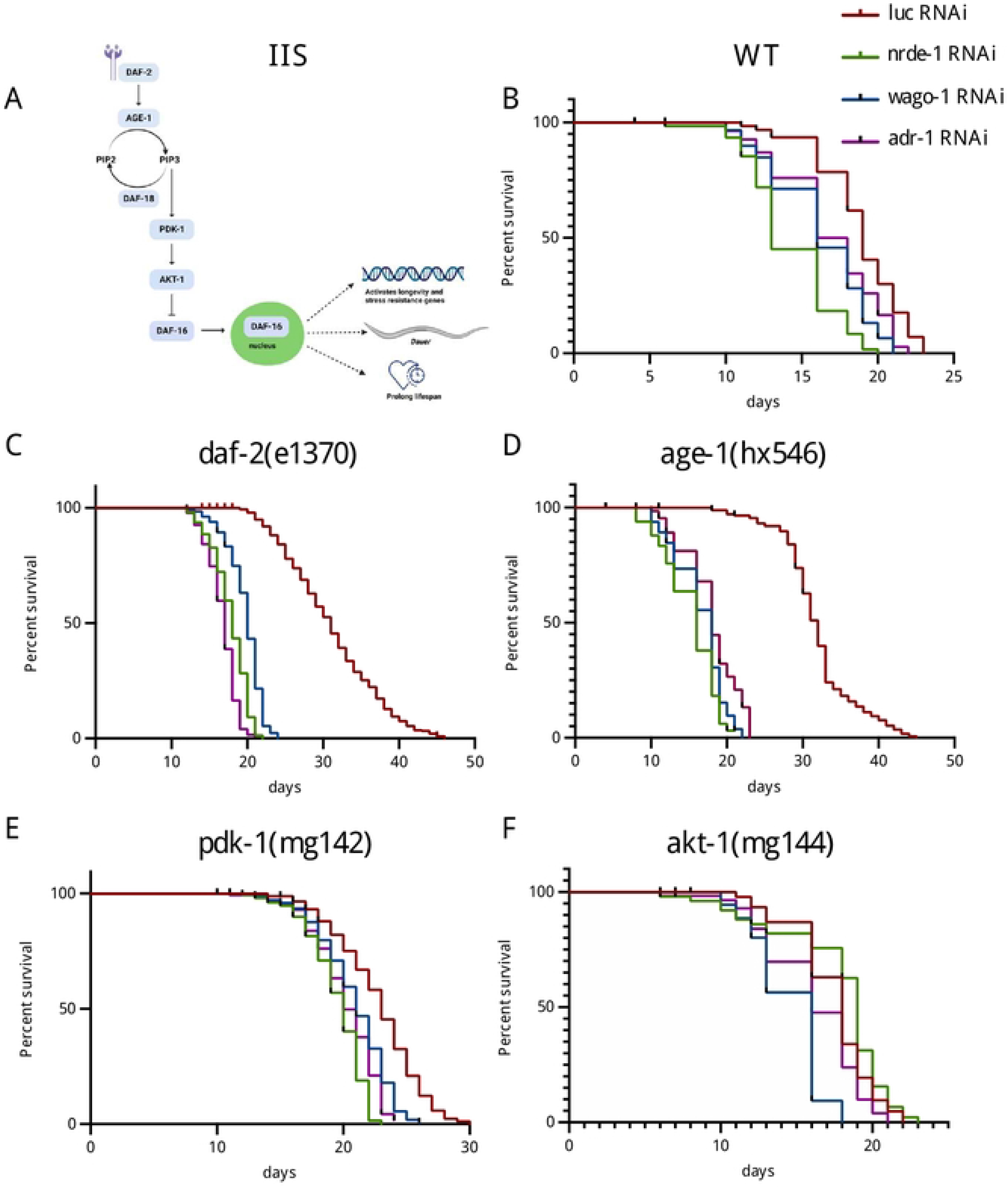
Insulin/IGF-1 receptor signalling (IIS)-mediated longevity is inhibited by *nrde-1*, *wago-1* and *adr-1* RNAi. A, the IIS pathway. B-F, Lifespan analysis representative of two independent experiments with similar results. *nrde-1, wago-1*, and *adr-1* were silenced by RNAi and RNAi targeting luciferase (luc) was used as a control. (B) N2 (WT), (C), *daf-2(e1370)*, (D) *age-1(hx546)*, (E) *pdk-1(mg142)* and (F) *akt-1(mg144)*. Data were compared using the log-rank (Mantel-Cox) test. Statistics are available in table S4.

### Other members of the nuclear RNAi pathway also control DAF-16 translocation, *dauer* formation, and lifespan

Given that the most pronounced effects obtained so far were with the *nrde-1* RNAi, we decided to test whether other members of the NRDE (*nuclear RNAi defective*) pathway were also involved in DAF-16 translocation. For that, we silenced *nrde-1, nrde-2, nrde-3* and *nrde-4* in *daf-2(e1370)* mutant worms expressing the DAF-16::GFP reporter construct or in heat-shocked WT worms expressing the same transgene. We then monitored DAF-16::GFP nuclear localization (Figure 4A-B). We observed that all members of the NRDE pathway inhibited DAF-16::GFP nuclear localization to some extent, although the effects were not as marked as those observed with *nrde-1* RNAi. Still, *nrde-4* RNAi resulted in a more pronounced inhibition of DAF-16 nuclear localization than *nrde-2* or *nrde-3* RNAi. These results demonstrate that the nuclear RNAi pathway controls DAF-16 function.

**Figure 4.**
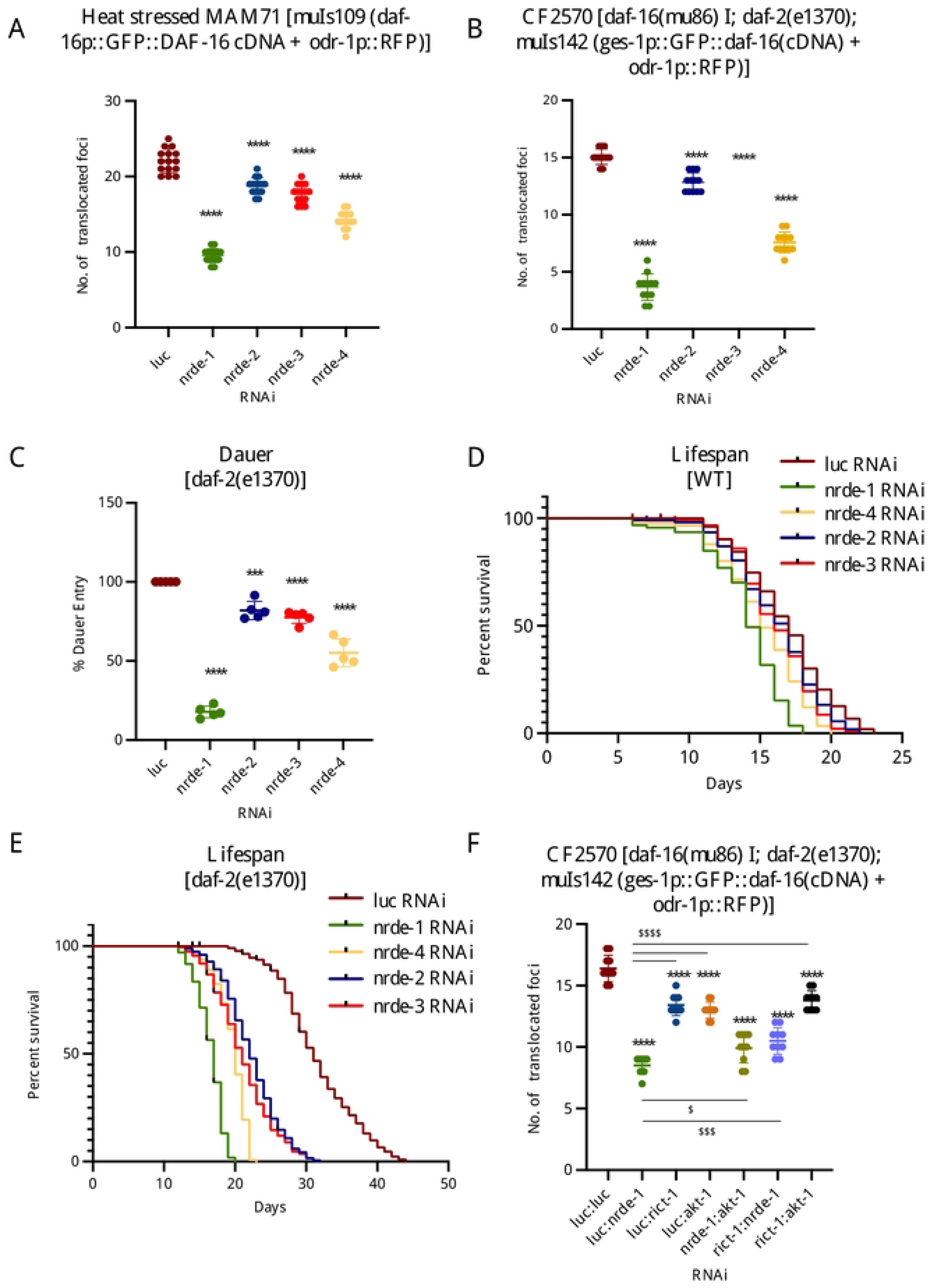
Other NRDE members also control DAF-16 function. A-E, *nrde-1, nrde-2, nrde-3*, and *nrde-4* were silenced by RNAi and RNAi targeting luciferase (luc) was used as a control. A-B, DAF-16 translocation assay. In A, MAM71 worms were maintained at 20 °C and heat-shocked at 34 °C for 1.5 hours on the first day of adulthood (n = 10-15 worms per group). In B, CF2570 worms were maintained at 15 °C till L2 and then transferred to 25 °C until adulthood (n = 10-15 worms per group). C, *Dauer* assay on *daf-2(e1370)* mutants (n = 100 worms per replicate in 5 replicates per group). \*\*\**P* < 0.001; \*\*\*\**P* < 0.0001 vs. luc. Two-tailed student *t*-test was used for individual comparisons against the control group (*luciferase*). These experiments were repeated 3 times with similar results. D-E, Lifespan analysis in N2 (WT) and *daf-2(e1370)* mutants, respectively. Representative of two independent experiments with similar results. Data was compared using the log-rank (Mantel-Cox) test. Statistics are available in table S5. F, A combination of *luc:nrde-1*, *luc:rict-1*, *luc:akt-1*, *nrde-1:akt-1*, *rict-1:nrde-1* or *rict-1:akt-1* RNAi were used and RNAi targeting luciferase (*luc:luc*) was used as a control. DAF-16 translocation assay in CF2570 worms. Worms were maintained at 15 °C till L2 and then transferred to 25 °C until adulthood (n = 10 worms per group, repeated 3 times with similar results). \*\*\*\**P* < 0.0001 vs. *luc:luc* (control) or ^$^*P* < 0.05, ^$$$^*P* < 0.001, and ^$$$$^*P* < 0.0001 vs. *luc:nrde-1* RNAi using One-Way ANOVA and Tukey’s multiple comparison as post-hoc test.

We also tested the role of NRDE members in *dauer* formation using *daf-2(e1370)* mutant worms (Figure 4C) and lifespan regulation using WT (Figure 4D) and *daf-2(e1370)* mutants (Figure 4E). Consistent with DAF-16 nuclear localization, silencing of the NRDE genes, particularly *nrde-1* and *-4*, resulted in a decrease in the *dauer* population. We also observed a reduction in the lifespan of WT and *daf-2(e1370)* mutants when other *nrde* genes were silenced, mainly *nrde-1* and *nrde-4*. These results confirm the role of nuclear RNAi in DAF-16-mediated *dauer* formation and longevity regulation.

### Downregulation of *nrde-1* does not lead to de-suppression of IIS genes

Given the role of nuclear RNAi in genome silencing, we hypothesized that NRDE-1 might be acting through the suppression of genes involved in the IIS pathway. If this hypothesis was correct, inhibition of *nrde-1* would result in upregulation of IIS genes which could contribute to hyperactivation of the pathway independent of the ligand. To assess that, we measured the expression of the members of the IIS pathway *age-1, daf-18, pdk-1*, and *akt-1* by RT-qPCR in WT and *daf-2(e1370)* mutants exposed to *nrde-1* or control RNAi (Figure S2A-D). Here we also used *daf-16* RNAi to assess the extent to which these genes are regulated by DAF-16 itself. It is interesting to note that gene expression of components of the IIS increased by many folds in the *daf-2(e1370)* mutant, which is consistent with a compensatory overexpression when the system is inactive. However, this upregulation did not appear to be mediated by DAF-16, as it is not inhibited by *daf-16* RNAi. Overall, *nrde-1* RNAi did not affect IIS gene expression in WT or *daf-2* mutant worms, except for an upregulation of *daf-18* in *daf-2(e1370)* mutants. Upregulation of *daf-18* could not account for the suppression of DAF-16 translocation in *nrde-1* RNAi-treated worms since DAF-18/PTEN acts as an inhibitor of the IIS pathway, counteracting AGE-1/PI3K action and promoting DAF-16/FOXO function (Ogg & Ruvkun 1998). Instead, *daf-18* upregulation could represent an attempt to counterbalance a hyperstimulation of the IIS pathway downstream of AGE-1 resulting from *nrde-1* inhibition. Hence, these results suggest that *nrde-1* RNAi does not inhibit DAF-16 translocation by de-suppressing the expression of upstream components of the IIS pathway.

### NRDE-1 acts partially through mTORC2-AKT-1 to control DAF-16 nuclear localization

Given these results and the fact that *nrde-1* RNAi not only suppresses *age-1* mutant-induced longevity but also reduces it when compared to the WT worms, we concluded that NRDE-1 controls DAF-16 function downstream of AGE-1/PI3K. Moreover, since *akt-1* mutation abrogates the capacity of *nrde-1* RNAi to reduce the lifespan in WT worms, we hypothesized that NRDE-1 is acting through AKT-1. We thought that somehow *nrde-1* inhibition would result in induction of AKT-1 activity independent of DAF-2/AGE-1 signalling. AKT can be phosphorylated and activated by PDK-1 or by mTORC2 [85]. Since *nrde-1* RNAi also reduces lifespan in *pdk-1(mg142)* mutants, we decided to test whether *nrde-1* acted via mTORC2 instead. *rict-1* encodes the *C. elegans* ortholog of mammalian Rictor, a key component of mTORC2. In *C. elegans*, *rict-1* is involved in the regulation of fat metabolism, nutrition, growth, and lifespan (Soukas et al. 2009). We, therefore, tested whether *nrde-1* RNAi requires *rict-1* to reduce DAF-16 translocation. To do this we used a double-RNAi feeding scheme (*e.g.*, knockdown of *akt-1* and *nrde-1, rict-1* and *nrde-1*, or *rict-1* and *akt-1*) and checked the DAF-16::GFP nuclear localization in *daf-2* mutant worms (Figure 4F). As expected, in the case of single *nrde-1* knockdown (*luc:nrde-1*), we confirmed the downregulation of DAF-16::GFP nuclear localization. A mild downregulation was observed in *luc:rict-1*, *luc:akt-1*, and *rict-1:akt-1* when compared to the control (*luc:luc*), suggesting a compensatory stimulus to reduce DAF-16 nuclear translocation when AKT-1 or mTORC2 are abrogated in the context of *daf-2* loss of function. However, silencing *akt-1* or *rict-1* partially reversed the *nrde-1* RNAi phenotype, from which it becomes evident that *nrde-1* interacts genetically with the mTORC2 pathway and *akt-1* to control DAF-16 translocation.

## DISCUSSION

Our main objective in this study was to determine to what extent DAF-16 activity could be impacted by epigenetic modulators such as chromatin remodelling proteins and proteins controlling small RNAs activity. Surprisingly, we discovered from the initial candidate screen that only a small subset of the examined genes had an impact on DAF-16 nuclear localization. Since *nrde-1*, *wago-1*, and *adr-1* RNAi exhibited more consistent effects on DAF-16 subcellular localization, we decided to focus on them from that point on. Importantly, we found that these genes had an impact on DAF-16 endogenous function. Among the DAF-16 target genes inhibited by *nrde-1* RNAi, for instance, mitochondrial superoxide dismutase (*sod-3*) acts as an antioxidant (Van Raamsdonk & Hekimi 2009), while *scl-1* encodes a putative secretory protein that is required for lifespan extension in *daf-2* and *age-1* mutants (Ookuma et al. 2003). We also demonstrate that *nrde-1*, *wago-1*, and *adr-1* are partially required for IIS-regulated phenotypes, such as *dauer* entry and longevity.

In an attempt to pursue the pathway through which NRDE-1 regulates DAF-16 function, we found that the short lifespan phenotype elicited by *nrde-1* RNAi requires functional AKT-1. Moreover, *akt-1* RNAi partially reverses the inhibition of DAF-16 nuclear translocation in *daf-2* mutants with reduced *nrde-1* expression. A similar effect is observed when the mTORC2 component *rict-1* is silenced. Taking these data together with our findings that NRDE-1 acts downstream of AGE-1, we propose a model where *nrde-1* RNAi induces mTORC2, which in turn leads to AKT phosphorylation and subsequent inhibition of DAF-16 nuclear translocation independent of AGE-1-mediated signalling. Noteworthy, *akt-1* and *rict-1* RNAi only partially reversed the effects of *nrde-1* RNAi on DAF-16 translocation. This could be due to the limited efficiency of the RNAi or to an alternative pathway acting in concert, although when we used the *akt-1(mg144)* mutant, no reduction in lifespan was observed in *nrde-1* RNAi-treated worms. The latter results support AKT-1 as a key player in *nrde-1*-mediated lifespan regulation.

We also show that other NRDEs have interactions with the IIS pathway at the level of DAF-16::GFP translocation, *dauer* formation, and lifespan regulation. NRDE-1 has dual associations in the nuclear RNAi pathway: one is to act in association with the other NRDE components on nascent pre-mRNA transcripts at the site of RNA Pol II activity and the other is to act closer to the site of histone H3 lysine 9 methylation (H3K9me) silencing at the chromatin level (Burkhart et al. 2011). NRDE-2 is a nuclear-localized, evolutionarily conserved protein that functions in association with NRDE-3 for the transport of siRNAs from the cytosol to the nucleus. It is recruited by NRDE-3/siRNA complexes to the RNAi-targeted nascent transcripts within the nucleus (Guang et al. 2010). Finally, NRDE-4 is a nematode-specific protein also described as necessary for the nuclear RNAi pathway (Burkhart et al. 2011). Curiously, we noticed that the magnitude of the effects of *nrde-2* and *-3* RNAi was less so than of *nrde-1* and −*4*. A plausible possibility is that different RNAis could produce different levels of gene expression inhibition. However, while NRDE-2 and −3 are necessary for NRDE-1 association with pre-mRNA and chromatin, it was observed that NRDE-4 is required for the association with chromatin only, promoting H3K9me (Burkhart et al. 2011). Supporting the idea that H3K9me levels might be somehow regulating DAF-16 nuclear localization and longevity, the inhibition of SET-25 – a histone methyltransferase, responsible for methylating the lysine of histone H3 in *C. elegans* (Towbin et al. 2012) – mimics the phenotypes observed by the inhibition of NRDE-1 and NRDE-4 (Table S1 and S2). Our work also supports previous data from the literature, where it was shown that the nuclear RNAi pathway can genetically interact with DAF-16/FOXO promoting germ cell immortality (Simon et al. 2018). Based on our findings, we anticipate that ADR-1 and WAGO-1 might also play a role in germ cell phenotype in *C. elegans*.

Our data also implicate RNA metabolism in DAF-16 function. ADR-1 and ADR-2 are the two isoforms of adenosine deaminases (ADARs) in *C. elegans*. ADARs, commonly known as RNA-editing enzymes, deaminate adenosines to create inosines in double-stranded RNA. *adr-1* is mostly expressed in the nervous system (*e.g.*, sensory neurons and cilia, the ventral nerve cord, motor neurons, and interneurons), the embryos, and the developing vulva of the *C. elegans*. It is required for normal chemotaxis, and vulval development and to prevent the silencing of transgenes in somatic tissues by RNAi (Tonkin et al. 2002; Knight & Bass 2002). ADAR loss of function mutations in *C. elegans* render worms short-lived, which is consistent with ADR-1 positively regulating DAF-16 (Sebastiani et al. 2009). We did not observe the effects of ADR-2 on DAF-16 nuclear localization in our initial screen. ADR-1 is a much larger protein than ADR-2 (964 and 495 residues, respectively), with an additional double-stranded RNA-binding motif, which might account for differences in cellular function (Tonkin et al. 2002). Accordingly, it was observed that ADR-1 and ADR-2 could have distinct roles in *C. elegans*, as in the case of vulva development (Tonkin et al. 2002). The action of ADR-1 may be directly in the edition of the *daf-16* mRNA, since it was observed in flies that *Foxo* mRNA can be edited by Adar (Khan et al. 2020). Interestingly, AKT1 can physically interact and phosphorylate ADARs in human cell culture (Bavelloni et al. 2019), suggesting that the interplay between the RNA editing pathway and DAF-16/FOXO function might have been conserved throughout evolution.

Finally, we also identified WAGO-1 as a regulator of DAF-16 function in *C. elegans*. WAGO-1 is an ortholog of members of the human Argonaute/PIWI family and is localized to the P granule and expressed in the germline (Shirayama et al. 2014). WAGO-1 is involved in the endogenous small interfering RNA (endo-siRNA) pathway and interacts with secondary 22G-RNAs. In the germline, it functions in a genome surveillance system and silences transposons, pseudogenes, aberrant transcripts, and regulates gene expression (Gu et al. 2009b). A role for WAGO-1-associated small RNAs in promoting H3K27 methylation was previously identified in *C. elegans* (Mao et al. 2015). That is potentially interesting because the inhibition of two lysine H3K27 demethylases, *jmjd-3.2* and *utx-1* extends lifespan in a *daf-16*-dependent manner. Thus, it is possible that the role of WAGO-1 in maintaining H3K27 methylation levels also impacts DAF-16 function and longevity (Guillermo et al. 2021).

In summary, our study unveils the role for different proteins involved with epigenetic regulation in *C. elegans* DAF-16 activity, thus playing a role in development and survival. Exercise and caloric restriction are two anti-aging strategies that promote changes in the epigenome (Horvath & Raj 2018; Li et al. 2011; Guerra et al. 2019). Indeed, the epigenome is intimately associated with biological age and epigenetic reprogramming has been envisioned as a way to reverse aging (Horvath & Raj 2018; Ocampo et al. 2016). Therefore, understanding how proteins involved in chromatin function and RNA metabolism control evolutionarily conserved longevity pathways may help in the development of new approaches to overcome aging complications.

## EXPERIMENTAL METHODS

### *C. elegans* stock maintenance and strains

The strains used were: N2 (wild-type), CB1370 *daf-2 (e1370) III*, MAM71 *muIs109 (daf-16p::GFP::DAF-16 cDNA + odr-1p::RFP)*, CF2570 *[daf-16(mu86) I; daf-2(e1370) III; muIs142 (ges-1p::GFP::daf-16(cDNA) + odr-1p::RFP)]*, TJ356 *zIs356 [daf-16p::daf-16a/b::GFP + rol-6(su1006)] IV*, CF1407 *[daf-16(mu86) I; muIs71 ((pKL99) daf-16ap::GFP::daf-16a(bKO)) + rol-6(su1006)]*, GR1310 *akt-1(mg144) V*, GR1318 *pdk-1(mg142) X*, TJ1052 *age-1(hx546) II*, CF1553 *muIs84 [(pAD76) sod-3p::GFP + rol-6(su1006)].* The *C. elegans* strains used in the experiments were maintained at 15-20°C in Petri dishes with NGM medium containing OP50-1 *E. coli* bacterial layer, except for the experiments involving RNAi where the HT115 (DE3) bacteria were used. OP50 plates were supplemented with 100 μg/mL streptomycin. RNAi plates were supplemented with 1 mM IPTG, 100 μg/mL tetracycline and 100 μg/mL ampicillin.

### RNAi

The RNAi assays were performed by feeding as previously described (Kamath et al. 2001). All RNAi clones were obtained from the Ahringer’s or Vidal’s RNAi libraries (Fraser et al. 2000; Rual et al. 2004). For the silencing of *adr-1, nrde-1, wago-1* and *set-25* genes, worms were synchronised and the eggs obtained were plated on RNAi [genes] or RNAi control [luciferase] plates. For all the experiments worms were developed in RNAi from L1 stage.

### Lifespan assay

Approximately 80-100 age-synchronized worms were monitored daily from day 0 of adulthood. The lifespan assays were performed at 20°C, except for experiments including *daf-2* mutants which were grown at 15°C until L2 to prevent *dauer* entry and then transferred to 25°C. Lifespan plates contained antibiotics, 0.25 μg/mL of amphotericin B to prevent fungal contamination and 0.5 μg/mL FUdR, an inhibitor of mitosis, to prevent offspring development and overpopulation.

### Gene expression

The gene expression was performed according to the (Bratic I, 2009) protocol. Pools of 100 3-day adult worms were transferred to 1.5 ml tubes and the RNA extraction was performed following the Trizol^®^ Reagent (Invitrogen) manufacturer’s instructions. Briefly, after the trituration of the sample in the presence of Trizol, 1/5 (v / v) chloroform was added to the tube. After 10 minutes standing at room temperature the samples were centrifuged in a refrigerated Eppendorf centrifuge at 12,000 × g for 15 min at 4°C. After centrifugation, the colourless phase of the tube was transferred to a new tube for the precipitation of RNA in the presence of isopropyl alcohol. After another centrifugation at 12,000 × g, 10 min, 4°C, a precipitate containing RNA could be identified. Ethanol 75% was added to wash the precipitate and then the ethanol removed and sterile water was added. The RNA was quantified in NanoDrop 2000c spectrophotometer. To perform the reverse transcription, we used 0.2-1 μg of total RNA. The reaction was conducted using the High-Capacity cDNA Reverse Transcription kit (Applied Biosystems) according to the manufacturer’s protocol. After the synthesis of the cDNAs, we quantified the expression of genes by real-time PCR. The reaction was conducted using the Maxima SYBR Green Master Mix (Fermentas), primers at the final concentration of 250 nM, and fluorescence was detected using the Applied Biosystems 7500 Real-Time PCR System. The amplification protocol used was: 50 °C for 2 min, 95 °C for 10 min and 40 cycles of 95 °C for 15 s, 60 °C for 20 s and 72 °C for 30 s. *his-10* was used as endogenous control. Primer sequences are available in Table S6.

### GFP fluorescence and microscopy (*GFP reporter analyses)*

Age-synchronized *C. elegans* were maintained in OP50-1 or RNAi plates until adulthood. Worms of the selected age were picked into the wells of microtitre plates (flat bottom Greiner). 80 μL of M9 media (22 mM Na_2_HPO_4_, 22 mM KH_2_PO_4_, 85 mM NaCl, 1 mM MgSO_4_) was added into the wells, the worms were immobilized using 0.1% sodium azide and images were acquired using the Cytation V microscope (Biotek). A constant gain was set in the plate reader for each condition. Images were analysed using ImageJ and integrated density was quantified. For the nuclear localization assays, the frequency of nucleated DAF-16::GFP was counted based on the number of cells with nucleated versus cytoplasmic DAF-16::GFP (Pinto et al. 2018).

### *Dauer* Assay

*Dauer* assays were performed using the *daf-2(e1370)* strain. This mutant spontaneously enters in *dauer* when kept at 25°C. Synchronized eggs using the bleaching method were placed onto NGM plates and grown at 25°C. Worms were scored for the presence of *dauer* and non-*dauer* under the microscope after 60-65 hours.

### Statistical Analysis

*C. elegans* survival under various conditions and of various genotypes was monitored as described in the figures. For the statistical analysis of lifespan, we used Log-rank test (Mantel-Cox). For the other experiments we used the two-tailed student *t* test for comparisons between two groups, and one-way or two-way ANOVA tests with Tukey’s or Dunnett’s post-tests, respectively, for comparisons between more than two groups, depending on the number of independent variables. The differences were considered statistically significant when the *P* value was < 0.05. The software used was GraphPad Prism (Version 8).

## AUTHOR CONTRIBUTIONS

Evandro A. De-Souza performed the initial mini-screen experiment. Silas Pinto trained Sweta Sarmah. Sweta Sarmah performed all other experiments. Adam A. Antebi and Marcelo A. Mori obtained funding and supervised the project. Sweta Sarmah, Evandro A. De-Souza, and Marcelo A. Mori wrote the initial draft. All authors read, contributed to, and approved the final version of the manuscript.

## ACKNOWLEDGEMENTS

We thank Dr Elzira Saviani for technical support. We thank the *C. elegans* Genetics Center (CGC) at the University of Minnesota, which is funded by the NIH Office of Research Infrastructure Programs (P40 OD010440), for providing *C. elegans* strains. This work was supported in part by the Conselho Nacional de Desenvolvimento Científico e Tecnológico (CNPq) – grant number 310287/2018-9, Coordenação de Aperfeiçoamento de Pessoal de Nível Superior - Brasil (CAPES) - Finance Code 001 and 88881.143924/2017-01, and Fundação de Amparo à Pesquisa do Estado de São Paulo (FAPESP) - grant numbers 2019/25958-9 (M.A.M.), 2017/01184-9 (M.A.M.), 2018/07703-0 (S.S.), 16/02207-0 (E.A.S.), 14/10814-8 (E.A.S.), and 17/04377-2 (S. P.).

